# Mapping and Genetic Dissection of Novel Tar Spot Resistance QTL on Maize Chromosome 1

**DOI:** 10.64898/2026.03.05.709888

**Authors:** Raksha Singh, Charles F. Crane, Tilahun Mekonnen, Sujoung Shim, Darcy E. P. Telenko, Stephen B. Goodwin

**Affiliations:** Crop Production and Pest Control Research Unit, U.S. Department of Agriculture-Agricultural Research Service (USDA-ARS), West Lafayette, IN 47907-2054, U.S.A; Department of Botany and Plant Pathology, Purdue University, West Lafayette, IN 47907-2054, U.S.A; Biotechnology Research Center, Addis Ababa University, Institute of Advanced Science and Technology, Addis Ababa, Ethiopia

**Keywords:** chromosome 1, corn, disease, maize, *Phyllachora maydis*, QTL, quantitative trait loci, resistance, tar spot

## Abstract

Tar spot, caused by the obligately biotrophic fungus *Phyllachora maydis*, significantly threatens maize (*Zea mays* L.) production across the Americas. Host genetic resistance offers the most viable long-term management strategy. Building on observed differential tar spot susceptibility, we utilized 92 recombinant-inbred lines (RILs) from the Intermated B73 × Mo17 (IBM-94) population to characterize the genetic architecture of resistance. Phenotypic analysis of 92 RILs plus the highly susceptible parent Mo17 and the moderately resistant parent B73 confirmed stable differences in susceptibility, with B73 consistently demonstrating moderate resistance compared to Mo17. Analysis of variance revealed highly significant genetic variation within the population (F = 12.96; p < 0.001). A high Pearson correlation (r = 0.8706, p < 0.0001) and coefficient of determination (R^2^ = 0.7579) across environments indicated that 76% of the phenotypic variance is attributable to genetic factors. Linkage mapping identified a novel, consistent major QTL cluster on chromosome 1. This cluster comprises five regions (qTAR_1.1 through qTAR_1.5) exceeding the significance threshold (LOD 3.8) in both years. We identified 74 candidate genes including bZIP and RING/U-box proteins at significant SNP peaks. Additionally, gene annotations revealed a high concentration of wall-associated kinases and S-locus lectin protein kinases within the qTAR_1.4 and qTAR_1.5 regions, alongside potential defense-related transcription factors (MYB, bZIP, and C2H2 zinc fingers). These results provide a framework for high-resolution mapping and functional validation to accelerate the development of tar spot-resistant maize cultivars.

---

Maize (*Zea mays L.*) is the world’s most widely distributed staple crop and a cornerstone of global food security. It serves as a versatile resource for direct human consumption, livestock feed, and bioenergy production (Godfray et al. 2010). However, this vital resource faces an escalating threat from various pathogens, among which is tar spot, an aggressive foliar disease that has rapidly transitioned from tropical curiosity to a premier concern for global agriculture. The disease is caused by the obligately biotrophic ascomycete *Phyllachora maydis*. It is characterized by the development of black, raised fungal structures called stromata on the leaf surface. These stromata are frequently surrounded by necrotic, tan halos, creating the diagnostic “fisheye” lesion. These lesions severely reduce the plant’s photosynthetic capacity and trigger premature senescence, ultimately leading to significant yield loss (Bajet et al. 1994; Mueller et al. 2020). Previous studies from Latin America attributed the fisheye phenotype to a synergistic complex involving *Monographella* (now renamed *Microdochium*) *maydis* and *P. maydis*; however, contemporary observations in the United States have not confirmed the presence of *M. maydis* (McCoy et al. 2019; Singh et al. 2025). Instead, current research suggests these necrotic symptoms may result from interactions between *P. maydis* and secondary opportunistic fungi, such as *Fusarium* spp. or *Paraphaeosphaeria neglecta* (Luis et al. 2023; McCoy et al. 2019). Although *M. maydis* has been reported recently at high frequencies in some parts of Ecuador and Guatemala (Sic-Hernandez et al., 2026) its role in tar spot etiology remains unknown.

Since its initial detection in Indiana and Illinois in 2015, *P. maydis* has spread with remarkable speed across the Midwest U.S. and into parts of southern Canada (Rocco da Silva et al. 2021). The pathogen overwinters effectively in crop debris, allowing inoculum to build up over successive seasons, especially in fields utilizing minimal tillage or continuous corn rotations (Groves et al. 2020). Under favorable environmental conditions, specifically moderate temperatures (18–23°C) and high leaf moisture, yield losses in susceptible hybrids can exceed 50%, representing a significant economic burden to the agricultural sector (Mueller et al. 2020; Webster et al. 2023). While fungicides offer temporary relief, the deployment of host resistance remains the most sustainable management strategy. Many commercial maize hybrids exhibit no or only partial resistance, leaving a critical need for more robust genetic resources, particularly within North American germplasm (Singh et al. 2018; Telenko et al. 2019). Efforts to address this gap have focused on screening broad-based maize germplasm, specifically targeting North American accessions to isolate traits that can be introgressed quickly into high-yielding, resistant cultivars (Cao et al. 2017; Lipps et al. 2022; Singh et al. 2023).

Previously, genetic studies using The International Maize and Wheat Improvement Center (CIMMYT) germplasm suggested that resistance might be controlled by major dominant genes; however, recent quantitative analyses indicate a more complex, polygenic architecture (Ceballos and Deutsch 1992; Hernandez-Ramos et al. 2015). Recent genome-wide association studies (GWAS) and linkage mapping have identified several quantitative trait loci (QTL), most notably qRtsc8-1 on chromosome 8, which has been validated across diverse African and tropical maize populations (Mahuku et al. 2016; Cao et al. 2017). Despite these advances, the efficacy of these loci from tropical maize lines against North American *P. maydis* populations remains a critical knowledge gap.

To dissect the genetic basis of resistance with higher precision, it is essential to utilize populations with increased recombination frequencies. The Intermated B73 x Mo17 (IBM) recombinant-inbred line (IBM Syn5) population was generated by intermating a B73 X Mo17 F_2_ population for five generations prior to six or more generations of self fertilization (Lee et al. 2002; McMullen et al. 2009). Due to the accumulation of crossover events during intermating, this population exhibits a nearly four-fold increase in recombination relative to standard RILs. Consequently, the IBM Syn5 population provides superior resolution for constructing high-density genetic maps and performing fine-scale QTL and heterosis analyses (Lee et al. 2002). The utility of the IBM Syn5 population is well established in the genetic dissection of numerous traits including resistance to foliar pathogens of maize. It has been instrumental in identifying QTLs for resistance to Northern Leaf Blight (NLB), where researchers leveraged the increased recombination to pinpoint narrow genomic intervals for genes governing incubation period and lesion expansion (Chung et al. 2010). Similarly, studies on Southern Leaf Blight (SLB) utilized the IBM population to resolve resistance into multiple small-effect QTLs, providing insights into the additive nature of maize defense (Kump et al. 2011). Furthermore, the population has been used to map resistance to Gray Leaf Spot (GLS) and Common Rust, often revealing “defense hotspots” that confer broad-spectrum protection (Balint-Kurti et al. 2007).

Based on our previous data, Mo17 showed a highly susceptible phenotype compared to B73, which was moderately resistant in response to natural infection by *P. maydis* (Singh et al. 2023). This differential response guided the selection of the IBM-94 core subset Syn5 population for the current analysis. Given that B73 and Mo17 are foundational to the U.S. hybrid maize industry, characterizing the resistance alleles within this pedigree is critical for breeding strategies against tar spot. Hence, as a strategic starting point, the IBM-94 Syn5 core subset was selected for initial disease screening rather than the larger mapping population. This allowed for an efficient survey of the genetic diversity inherent in the B73 x Mo17 pedigree while optimizing field resources and labor. This approach served as a proof of concept to validate the heritability of tar spot resistance and to standardize phenotyping protocols under the specific environmental conditions of northern Indiana. By utilizing this representative subset, we confirmed that sufficient phenotypic variation existed to justify the potential scaling to a larger mapping population in future analyses. The primary objectives of this research were to: (i) evaluate the disease phenotypes of 94 IBM Syn5 lines and their parents B73 and Mo17 under natural infection by *P. maydis* across two years (2020 and 2021) in the field; (ii) test for stable QTLs governing tar spot resistance using linkage mapping with genome-wide, genotyping-by-sequencing (GBS)-derived single-nucleotide polymorphisms (SNPs); and (iii) identify candidate genes within the primary QTL confidence intervals. These findings establish a framework for the high-resolution mapping and functional validation of the tar spot resistance in maize inbred line B73.

## Materials and Methods

### Plant material and field trials

The Intermated B73 x Mo17 (IBM) recombinant-inbred line population, specifically the IBM-94 (Syn5) core subset, was utilized in this study. This population, consisting of 94 RILs plus the parental lines B73 and Mo17, was obtained from the Maize Genetics Cooperation Stock Center (Urbana, IL). The Syn5 population was developed by intermating B73 × Mo17 for five generations before more than six rounds of self fertilization, creating a high-resolution genetic resource with four times the recombination of standard RILs (Lee et al. 2002). Given that B73 displays moderate resistance while Mo17 is highly susceptible to *P. maydis* (Singh et al. 2023), the IBM-94 core subset RIL population provides a more efficient start for disease screening rather than the larger mapping population for dissecting the genetic basis of tar spot resistance.

The complete set of 94 RILs, along with the resistant parent B73 and susceptible parent Mo17, were planted in replicated field trials at the Pinney Purdue Agricultural Center (PPAC), Wanatah, IN. However, due to poor germination in both the 2020 and 2021 seasons, two lines (RILs #M0044 and #M0054) were excluded from the analysis, resulting in a final population of 92 RILs. The PPAC location was chosen due to its consistent history of high tar spot disease pressure in previous years. As *P. maydis* is an obligately biotrophic fungus and cannot be cultured for inoculation, all field trials relied on natural infection. Plots were planted on June 15, 2020, and May 27, 2021, respectively, for the first- and second-year trials. Each plot consisted of two rows, approximately 3.0 m (10 ft) wide and 9.1 m (30 ft) long, with an approximate planting density of 20 seeds per row. Each experiment was laid out with two replications. The soil type at this location is a Sebewa loam. At all plot locations, the previous crop was corn with a history of tar spot. Fungicide treatments were not applied during the two-year experimental period to allow for natural disease development.

### Phenotypic evaluation

Phenotypic evaluation of tar spot symptoms commenced after the onset of natural infection and was conducted as described previously (Singh et al. 2023). Disease scoring was performed at three stages of disease development: early (tassel vegetative growth; early September), middle (reproductive growth R2 [blister]; mid-September), and late (reproductive growth R5 [dent]; early October). On 1 September 2020, and 7 September 2021, when plants were at the tassel (VT) growth stage, the field trials were rated for early tar spot severity by visually assessing the percentage of symptomatic area per leaf following online training (https://severity.cropprotectionnetwork.org/crop/corn/tar-spot-stroma). This 0-to-100% disease severity scale at different growth stages has been used as a standard in numerous prior publications (Kleczewski et al. 2020; Telenko et al. 2019, 2021, 2022; Singh et al. 2023). The small, dark, raised stromata of tar spot are distinct from those of other corn diseases (Rocco da Silva et al. 2021), allowing for accurate scoring even on slightly senescent leaves. To ensure pathogen identity, several samples were collected at the beginning of the experiment, and presence of *P. maydis* was confirmed by sequencing the internal transcribed spacer region with standard procedures (data not shown). Four plants were rated from each line in two replicates, beginning 2 weeks after the first observed disease incidence. To capture whole-plant disease severity, ratings were taken from three parts of the canopy: lower leaf (defined as two leaves below the ear leaf), ear leaf, and upper leaf (defined as two leaves above the ear leaf). The means of these three ratings were used for calculating the disease severity for each plant. Mid-season disease severities were rated on September 15, 2020, and September 22, 2021, at the VT to silk (R1) growth stages. Late disease severity was rated on October 1, 2020, and October 2, 2021, at the R1 to physiological maturity (R6) growth stages. Disease scoring methods and field trial locations remained consistent across both years except for the plots.

### Statistical analysis of phenotypic data

Analysis of variance (ANOVA) was performed on phenotypic data for recombinant-inbred lines (RILs) and parental lines, considering individual observations across both growing seasons, using an R package. Significant differences between individuals were tested with Tukey’s test at a 5% significance value (α=0.05). A Pearson correlation analysis was performed to determine the correlation of disease severities between 2020 (Environment I) and 2021 (Environment II). Phenotypic data from the 2020 and 2021 field seasons were organized and analyzed using Microsoft Excel and R statistical software. To assess the repeatability of the tar spot disease screening, a Pearson correlation coefficient (r) and a coefficient of determination (R^2^) were calculated for the 92 IBM RILs that were successfully scored in the field across both environments. The statistical significance of the correlation was determined using a two-tailed t-test with n - 2 degrees of freedom to derive the p value. The standard error of the correlation (SE_r_) was calculated using the formula: SE_r_ = \sqrt{\frac{1 - r^2^} {n - 2}}. A linear regression model was applied to the data to visualize the relationship between years and to identify potential outliers. All hypotheses were tested at a significance level of α = 0.05.

### QTL detection by linkage mapping

Quantitative trait loci were identified using the scan1 and find_peaks functions within the R package qtl2 (Broman et al. 2019). Phenotypes used for mapping included replicated disease scores from 2020 (Environment I), 2021 (Environment II), or a combined average across both years. Genotype data were sourced from the VCF-formatted file “ZeaGBSv27_publicSamples_imputedV5_AGPv4-181023.vcf”, downloaded from Panzea (data.cyverse.org, directory /shared/panzea/genotypes/GBS/v27). This file contained 943,455 GBS-based SNP and small indel markers from 17,280 inbred lines, including 94 lines from the B73 × Mo17 NAM population. Parental genotypes (Mo17 and B73) were obtained from the hmp321_agpv4_chr[1-10].vcf files available at the specified URL. Previously missing genotypes in this dataset had been imputed with LinkImpute (Money et al. 2015). The downloaded VCF file underwent several processing steps using custom Perl scripts. First, a script (phasemarkers0929.pl) extracted genotypes of segregating markers in the 94 B73 × Mo17 inbred lines, ordering them by chromosome and nucleotide coordinate in the B73 maize genome, version 4. This script required parental allele frequencies to fall within the interval [0.3, 0.7]. Next, the segregating markers (last 112,202 lines of output) were copied to a file, and purgemissingmarkers.pl removed all markers with more than 5% missing data and heterozygotes, reducing the map to 30,377 markers. At this stage, the genotype file included marker names, genotypes, chromosomes, and coordinates.

The script ironoutphases.pl identified instances where the recombination fraction between consecutive markers exceeded 0.5, indicating a switch in parental phase. It interchanged allele designations for subsequent markers, resulting in a map with uniform parental phase across each chromosome. This process did not guarantee consistent parental phase designations between different chromosomes. Although this reduced the recombinational map length, it remained excessive due to cumulative sequencing errors. correctsingletons.pl attempted to correct sequencing errors where genotypes switched twice within a triplet of consecutive markers in a single mapping individual, further reducing the map length. Finally, makemap09302024.pl formatted the markers for use by R/qtl2. Recombination fractions were not transformed using either the Haldane or Kosambi mapping functions; the Haldane function would have further lengthened the already long map. The order of markers in the B73 assembly was presumed to be correct.

Once QTL peaks were identified with the qtl2 function find_peaks, the script pickmarkers1003.pl identified gene models within 10 or 15 cM of a QTL peak position in the linkage map. This generated a merged, ordered list of markers and gene models as candidates for further investigation. Gene models were downloaded from Phytozome (https://phytozome-next.jgi.doe.gov/info/Zmays_RefGen_V4) as Zmays_493_RefGen_V4.gene.gff3 and Zmays_493_RefGen_V4.annotation_info.txt. The proportion of phenotypic variance explained (PVE) was calculated from Logarithm of the Odds (LOD) peak values using the equation PVE = 1−10^(−2⋅LOD/n)^, where LOD represents a peak LOD score and n is the number of genotyped individuals (Broman and Sen. 2009). Permutation tests (1000 permutations) were performed using the scan1perm function in R/qtl2 (Broman et al. 2019) to determine p values for LOD thresholds at 1%, 5%, and 10%. Additive effects of each QTL on the phenotype were estimated with the scan1coef function in R/qtl2. Because RIL populations lack heterozygotes, it was not possible to estimate dominance effects. Significant QTL positions and associated SNP markers were shown using Mapchart (Voorrips 2002).

### Candidate gene annotation

Candidate genes were identified based on the significantly associated SNPs detected from the linkage mapping, utilizing the maize B73 reference genome V4 for GBS SNPs, available from the MaizeGDB website (https://phytozome-next.jgi.doe.gov/info/Zmays_RefGen_V4) via Zmays_493_RefGen_V4.gene.gff3 and Zmays_493_RefGen_V4.annotation_info.txt files. Putative candidate gene annotations were performed using the following databases: MaizeGDB, National Center for Biotechnology Information (https://www.ncbi.nlm.nih.gov/), Gramene (http://www.gramene.org/), and PlantGDB (http://www.plantgdb.org/ZmGDB/).

## Results

### Phenotypic variation of tar spot resistance in the maize Intermated B73 x Mo17 (IBM) recombinant-inbred line mapping population

Tar spot severity in the parental inbred lines B73 and Mo17, along with 92 Mo17 x B73 recombinant-Inbred Lines (RILs), was evaluated across two distinct growing seasons: the 2020 field (Environment I) and the 2021 field (Environment II). Seeds of two lines failed to germinate, so were excluded from further analyses. Although conducted at the same site, different field plots were used in both years which have variable sources of inoculum and pathogen populations. These factors, coupled with differences in temperature, rainfall, and humidity between the years, resulted in the classification of these as distinct environments (Supplementary Table S1). Disease severity was determined using a published protocol (Singh et al. 2023), confirming that B73 is moderately resistant and Mo17 is highly susceptible at 110 Days After Planting (DAP). Consistently across both environments (Supplementary Table S2), B73 showed significantly lower disease severity (Mean Disease Severity (MDS) of 8.3% in Environment I and 10.5% in Environment II) compared to Mo17 (MDS of 17.0% in Environment I and 33.4% in Environment II), demonstrating consistent differences in susceptibility. The frequency distribution of tar spot severity among the 92 RILs demonstrated environmental dependence (Fig. 1, A and B). In Environment I, the distribution exhibited a strong positive skew towards resistance (Fig. 1A). The majority of the population (69 RILs) displayed symptoms between 5 and 15% mean disease severity, suggesting a high degree of resistance within the population in this environment (Supplementary Table S2). Conversely, Environment II showed an irregular and likely multimodal distribution, indicating the coexistence of at least two phenotypic groups (Supplementary Table S2). The data were broadly spread (87 RILs between 10 and 35% severity), with distinct clusters evident in both the low-to-mid (10 to 20%) and the mid-to-high ranges (25 to 35%).

**Fig. 1.**
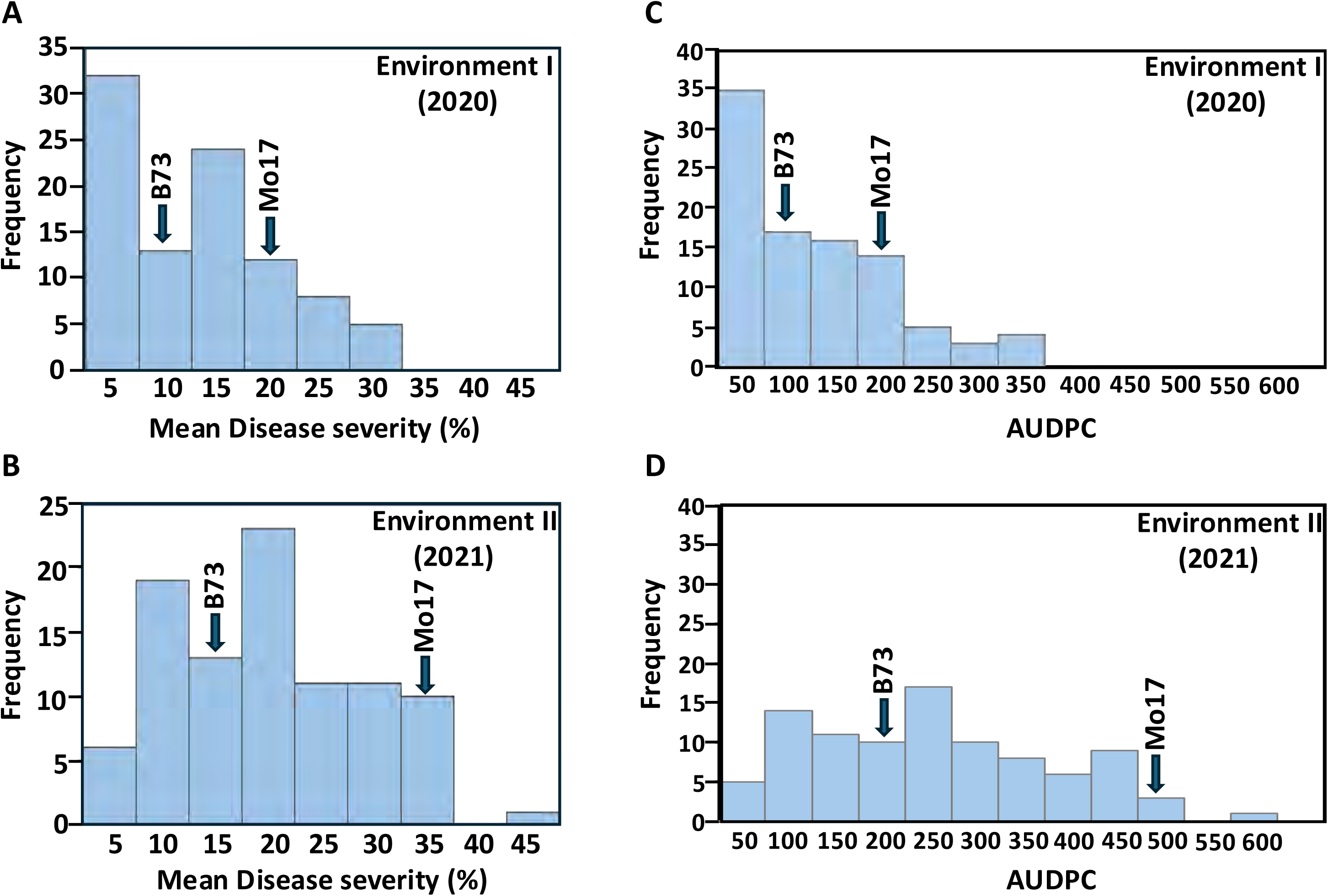
Frequency distribution of mean disease severity and Area Under the Disease Progress Curve (AUDPC) for tar spot disease in 92 recombinant-inbred lines from the Intermated B73 × Mo17 (IBM) segregating Syn5 population in field trials over across two environments, the 2020 (Environment I) and 2021 (Environment II). **A**, Mean disease severity (%) in Environment I (2020) and **B**, Environment II (2021). **C**, AUDPC scores in Environment I (2020) and **D**, Environment II (2021). Arrows indicate the mean performance of the parental lines B73 (moderately resistant) and Mo17 (highly susceptible). In all panels, the x axis represents the disease metric, and the y axis indicates the frequency of recombinant-inbred lines (RILs).

Frequency distributions for the Area Under Disease Progress Curve (AUDPC) similarly suggest a quantitative genetic control for tar spot resistance (Fig. 1, C and D). In 2020, the distribution was heavily skewed toward lower AUDPC values, with over 30 lines falling into the lowest bin (0–50). In contrast, the 2021 distribution was more broadly dispersed, indicating higher overall disease pressure and a shift toward increased susceptibility (Fig. 1.D). Pearson Coefficient correlation analysis between environments revealed a strong and highly significant positive correlation (r = 0.8706, p < 0.0001) between tar spot severity across the two-year (2020 and 2021) study period. The coefficient of determination (R^2^ = 0.7579) indicates that approximately 76% of the phenotypic variance was consistent across both years, suggesting robust genetic control over the evaluated resistance traits (Fig. 2). The reliability of the phenotyping protocol was further validated by the performance of standardized checks: resistant RILs (green) consistently maintained low severity, while susceptible lines (red) occupied the high-severity quadrant of the regression (Fig. 2). While individual outliers such as RIL# 3409-27 IBM RI M0043 exhibited seasonal variation, the overall population maintained high linearity (Supplementary Table S3). Furthermore, the pairwise Pearson correlation of AUDPC across both environments remained highly significant (r = 0.787, p<0.001) (Fig. 3). These findings confirm that the disease screening-methodology provides the necessary precision for reliable identification and selection of stably resistant germplasm.

**Fig. 2.**
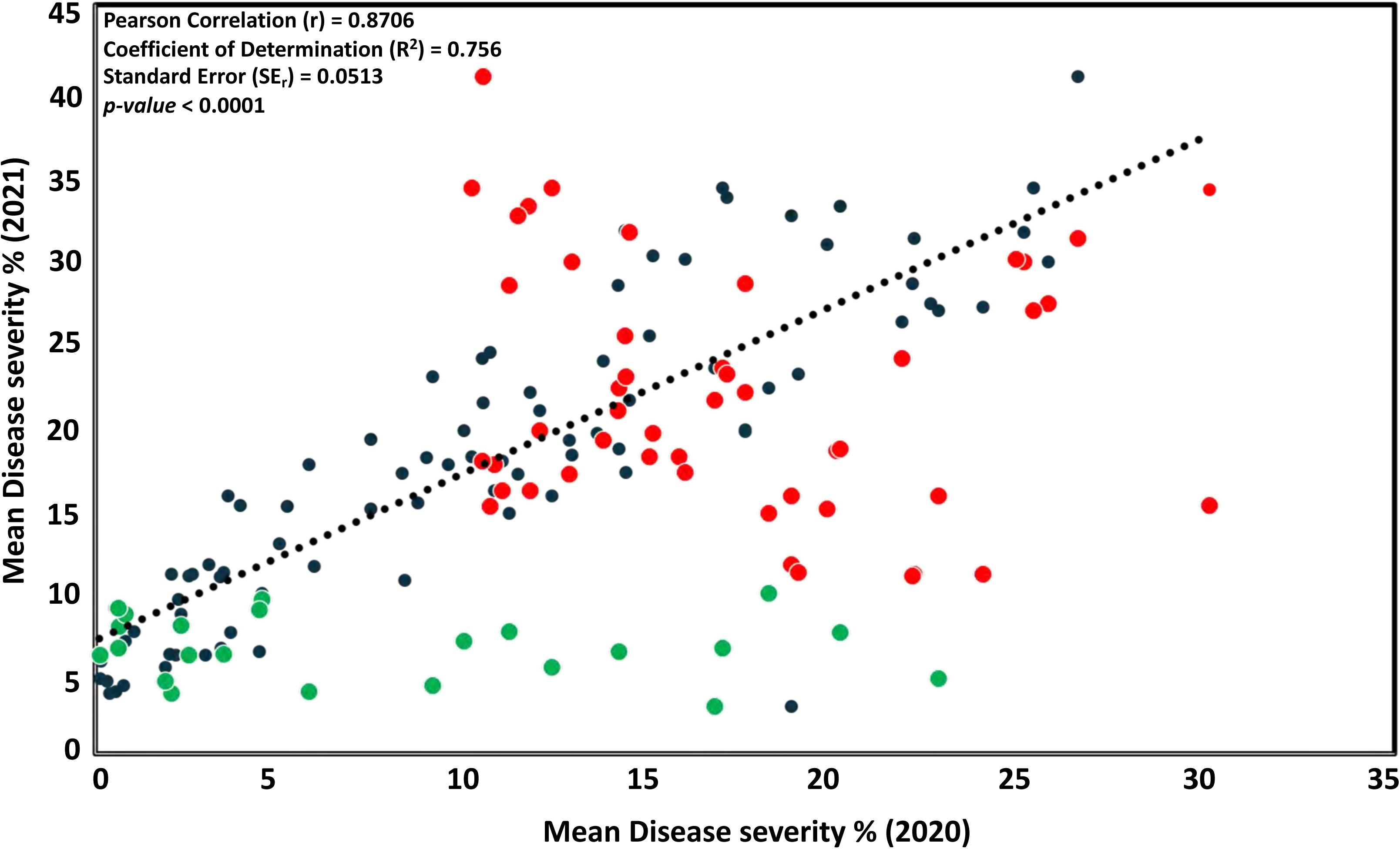
Correlation of tar spot mean disease severity index scores across two years (2020 and 2021) of field trials. Scatter plot illustrating the phenotypic relationship between disease severity percentages recorded in 2020 (x axis) and 2021 (y axis) for 92 Intermated B73 × Mo17 (IBM) maize recombinant-inbred lines. Dark blue circles represent the experimental population, while resistant and susceptible check inbreds are highlighted in green and red, respectively. The dashed line indicates the linear regression model (R^2^ = 0.7579). Correlation analysis revealed a strong, statistically significant relationship (r = 0.87, p <0.0001, t-statistic = 16.97) between disease phenotypes of the experimental population in the two years.

**Fig. 3.**
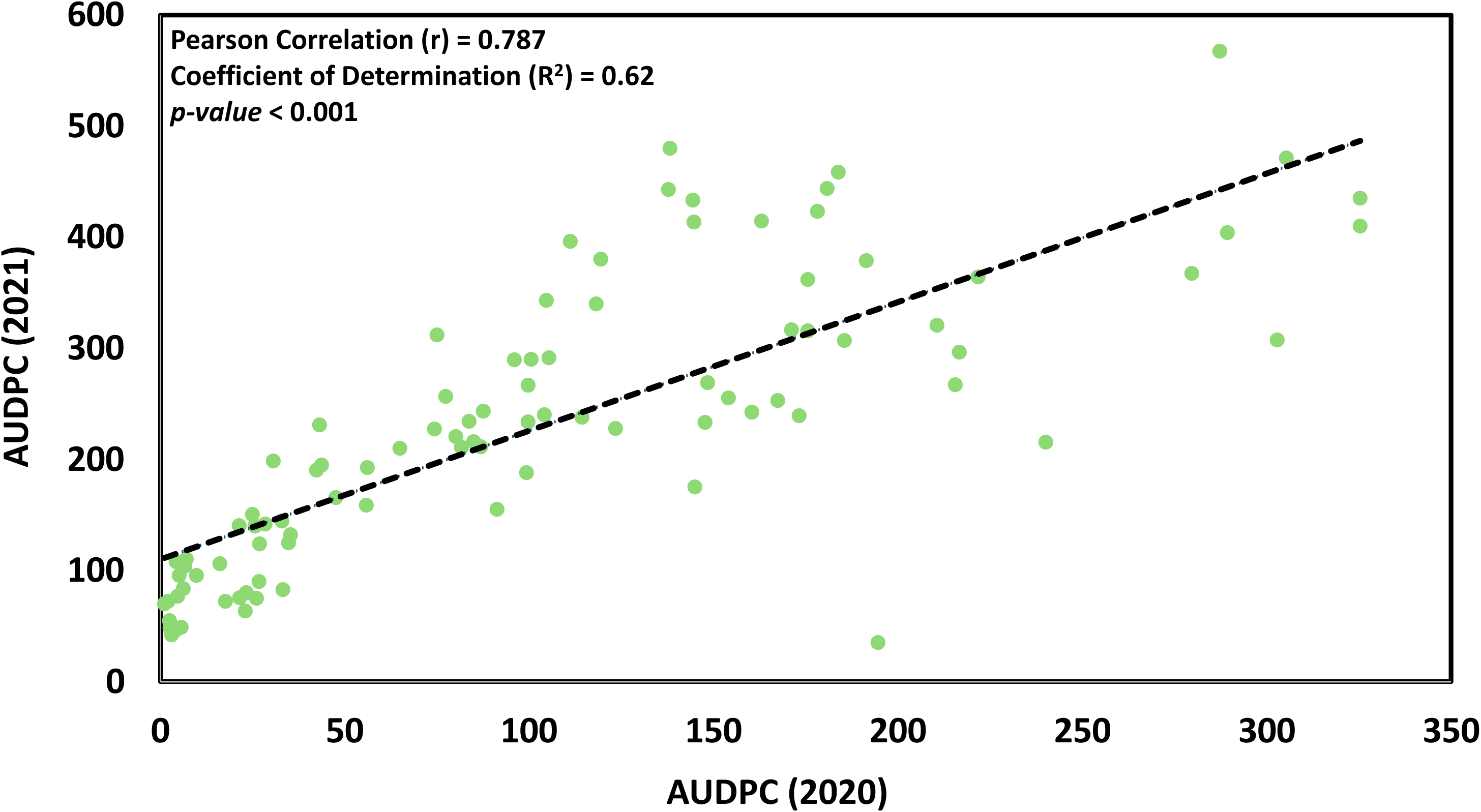
Correlation of tar spot severity measured by Area Under the Disease Progress Curve (AUDPC) between 2020 and 2021 across 92 Intermated B73 × Mo17 (IBM) maize lines. A strong positive correlation was observed (r = 0.787, p < 0.0001), indicating consistent genetic resistance across environments. Despite the correlation, a paired t test revealed significantly higher disease pressure in 2021 compared to 2020 [t(91) = -15.87, p < 0.001)].

There was a marked increase in tar spot severity from 2020 to 2021. The mean AUDPC increased more than twofold, rising from 99.67 to 218.43. A paired t-test confirmed that the difference in disease progression between the two environments was highly significant (*p < 0.001*) (Fig. 4, A and B). Box plots illustrate this disparity, showing a higher median and a wider interquartile range in 2021 compared to 2020, highlighting the segregation of diverse resistance phenotypes (Figure 4B).

**Fig. 4.**
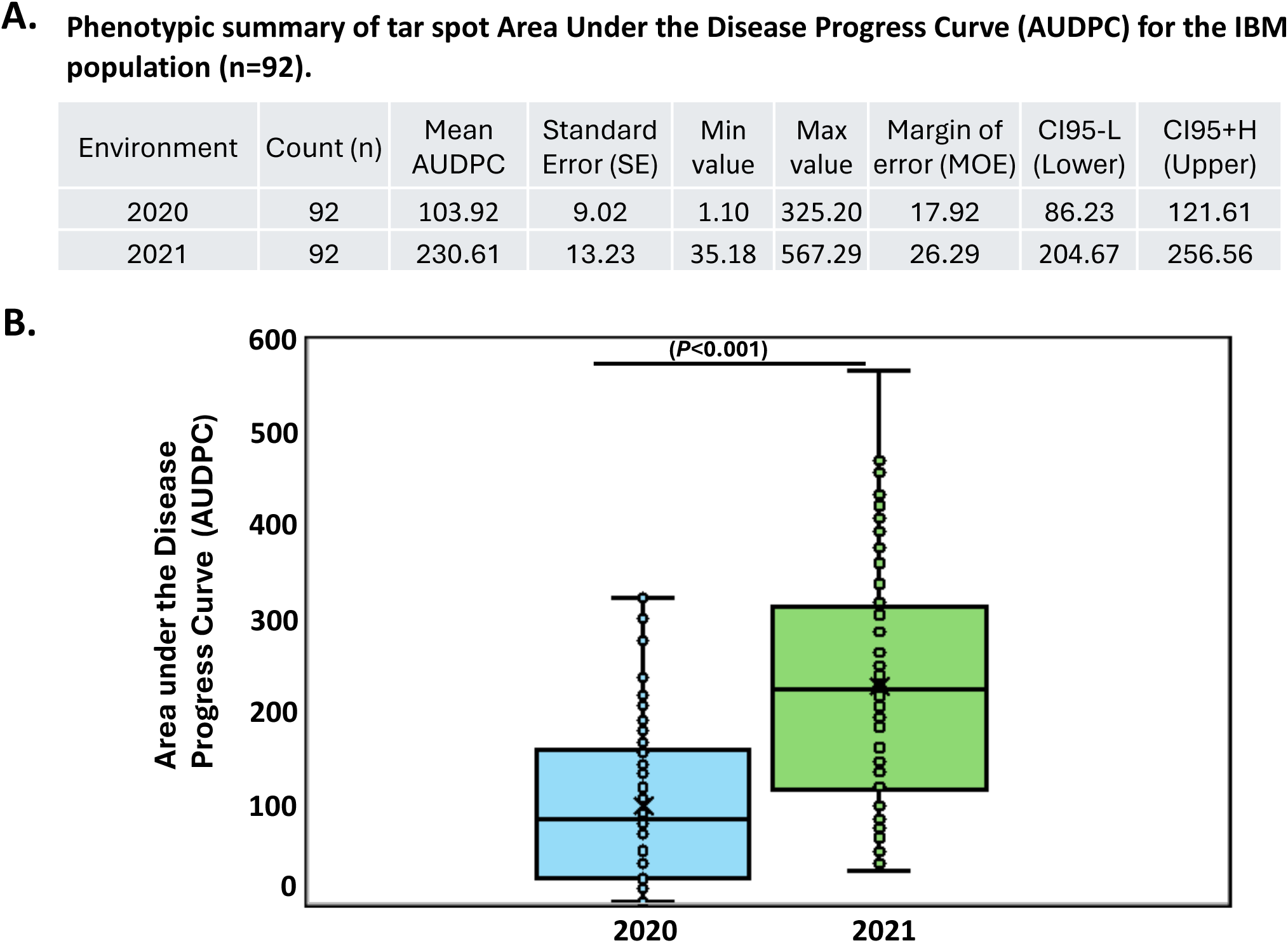
Phenotypic evaluation of tar spot resistance in the Intermated B73 × Mo17 (IBM) recombinant-inbred line (RIL) population (n = 92; two of the 94 RILs did not germinate) across two environments. **A**, Summary statistics for Area Under the Disease Progress Curve (AUDPC). **B**, Box plots showing median and interquartile ranges, with individual RIL means overlaid as jitter points. Significant difference between environments is indicated (P < 0.001, paired t-test).

The combined analysis of variance (ANOVA) for tar spot disease severity revealed highly significant difference among the 92 genotypes tested over two years at a single location (F = 12.96; *p<0.001*), indicating substantial genetic variation for resistance. The effect of replication also was significant (F = 211.27; *p<0.001*), reflecting environmental differences between the two years. In contrast, the variation among blocks within replication was not significant (F = 0.68; p = 0.72), suggesting that the experimental blocking effectively minimized within-year environmental variation (Table 1). The residual mean square was relatively low (Mean of Squares = 10.7), indicating good experimental precision. Collectively, these results demonstrate that selection for tar spot disease resistance among the evaluated genotypes is feasible and reliable, and that the observed differences are primarily due to genetic factors rather than experimental noise.

**Table 1.** Combined analysis of variance (ANOVA) for tar spot disease severity in 92 recombinant-inbred lines (RILs) from the Intermated B73 × Mo17 (IBM) Syn5 population (seeds from two of the 94 RILs failed to germinate) evaluated across two environments (Environment I in 2020 and Environment II in 2021) in Wanatah, Indiana.

| Source of variation | Degrees of freedom | Sum of squares | Mean square | F value | Pr(>F)<sup>a</sup> |
| --- | --- | --- | --- | --- | --- |
| Replication<sup>b</sup> | 1 | 2254 | 2253.7 | 211.27 | <2e-16<sup>***</sup> |
| Block(replication)<sup>c</sup> | 18 | 66 | 7.3 | 0.68 | 0.72 |
| Genotype | 93 | 12860 | 138.3 | 12.96 | <2e-16<sup>***</sup> |
| Residuals | 84 | 896 | 10.7 |  |  |
| Total | 196 |  |  |  |  |

<sup>a</sup> Replications within environments.
<sup>b</sup> Blocks nested within replications.
<sup>c</sup> \*\*\*= very highly significant at the $p < 0.001$ significance level.

### Quantitative trait locus (QTL) mapping for tar spot resistance

To dissect the genetic basis of tar spot resistance, linkage mapping was conducted using mean disease severity scores from 92 IBM Syn5 RILs across the 2020 and 2021 environments. A genome-wide significance threshold of LOD 3.8 (*p < 0.05*) was established based on 1,000 permutation tests to distinguish true genetic associations from background noise. QTL analysis was performed at early, mid, and late disease development stages. At the early stage, no definitive QTL surpassed the LOD 3.8 threshold. This lack of significant detection is likely attributable to incomplete disease progression, which limited phenotypic discrimination between genotypes at that time point (Supplementary Fig. S1). At the mid-disease stage, a QTL was identified on Chromosome 1, reaching a maximum LOD score of approximately 3.7 (Fig. 5A). Although this peak was the most prominent genomic feature, it remained slightly below the stringent significance threshold. In Environment II (2021), this same region on Chromosome 1 was again identified as a putative QTL with a LOD score of 2.9 (Fig. 5B). The consistency of this peak across both growing seasons, despite varying disease pressures, suggested that Chromosome 1 harbors a stable locus contributing to *P. maydis* resistance. By the late disease stage, the analysis revealed a consistent genetic architecture for resistance across both testing years (Fig. 6). A single major QTL was identified on chromosome 1 that significantly exceeded the genome-wide threshold in both 2020 and 2021, indicating a stable and primary genetic driver for tar spot resistance within this population. The high LOD score associated with this peak suggests it explains a substantial proportion of the phenotypic variance.

**Fig. 5.**
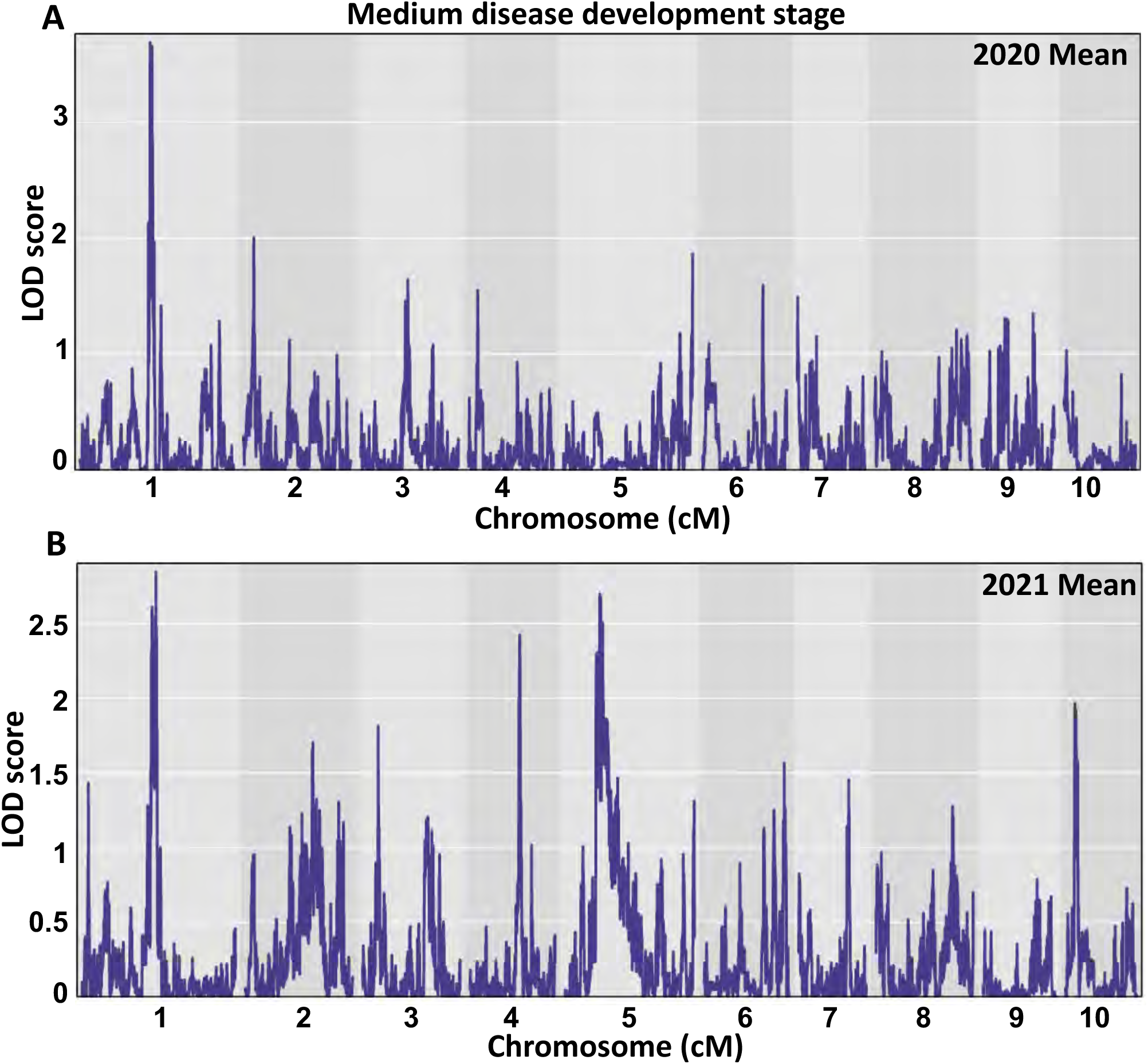
Genome-wide linkage analysis of tar spot resistance in the Intermated B73 × Mo17 (IBM) Syn5 maize recombinant-inbred line (RIL) population evaluated at the medium disease development stage across two years. Logarithm of the odds (LOD) profiles across 10 chromosomes are shown for 92 (two of the 94 lines failed to germinate) IBM RILs evaluated in the (**A**) 2020 and (**B**) 2021 field environments. The x axis represents the genetic position across all 10 chromosomes, and the y axis indicates the LOD score. A QTL on chromosome 1 was detected at the same genomics location in both years but was just below the significance threshold of LOD 3.8 (p < 0.05), determined through 1,000 permutations. In 2021, other minor, putative peaks were identified below the significance threshold on chromosomes 4, 5, and 10.

**Fig. 6.**
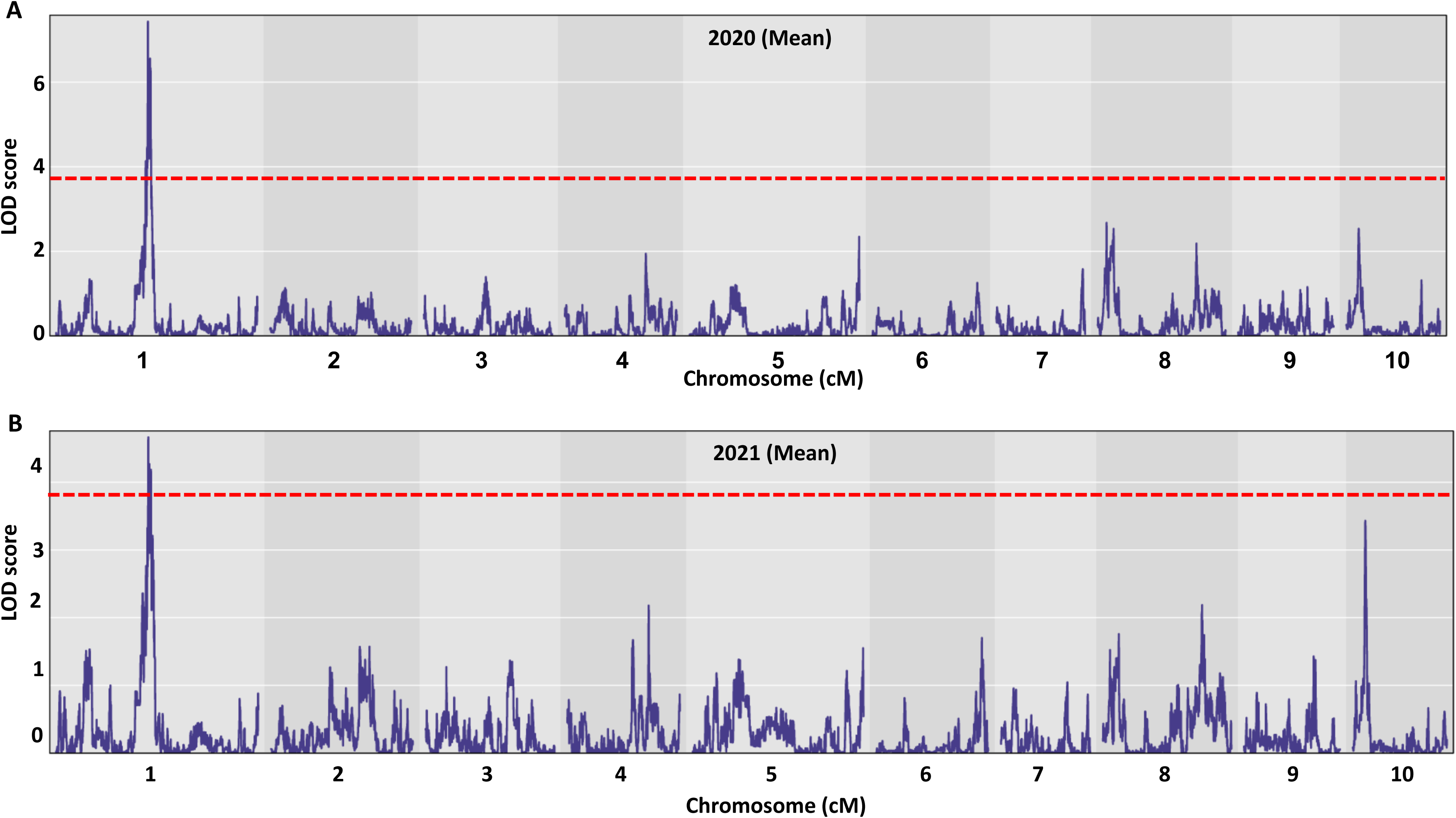
Genome-wide linkage analysis for tar spot resistance associated with quantitative trait loci (QTL) in the Intermated B73 × Mo17 (IBM) Syn5 maize recombinant-inbred line (RIL) population. The logarithm of the odds (LOD) score profiles over 10 maize chromosomes for 92 IBM RILs (seeds from two of the 94 RILs did not germinate) evaluated in the (**A**) 2020 and (**B**) 2021 field environments for the late disease development stage. The x axis represents the chromosomal positions based on Panzea genotypic data, and the y axis represents the LOD score indicating the measure of statistical significance of the association between a genetic marker and the trait. The horizontal, red, dashed lines denote the genome-wide significance threshold of LOD 3.8 (p <0.05), determined through 1,000 permutations. A major, consistent QTL exceeding the threshold was detected on chromosome 1 in both years and is provisionally designated *qRtsc1-1*. Additional minor QTL peaks exceeding a LOD score of 2.0 but below the overall threshold were observed in both years on chromosomes 8 and 10.

In addition to the major locus on chromosome 1, several minor QTL peaks were identified. Loci on chromosomes 8 and 10 exceeded a LOD score of 2.0 but remained below the rigorous genome-wide significance threshold of 3.8 (Fig. 6A and B). Notably, while previous studies on CIMMYT maize populations have detected major QTLs on chromosome 8, the corresponding peak for that QTL in this study was statistically insignificant, falling below the LOD 3.8 threshold.

Through linkage mapping, a total of 31 associated single-nucleotide polymorphisms (SNPs) were statistically significant and distributed across five adjacent QTL regions on chromosome 1: thirteen markers in the qTAR1.1 region; six markers in the qTAR1.2 region; three markers in the qTAR1.3 region; four markers in the qTAR1.4 region; and five markers in the qTAR1.5 region (Table 2). Several markers were located directly within peak regions (Fig. 7). For instance, S1_182626134 and S1_182627236 were in the qTAR_1.1 region (QTL position 1395.70 cM, QTL size 12.46 cM) and showed a Phenotypic Variation Explained (PVE) of 19.4% with an additive effect of -3.50 (Table 2), where negative additive effects indicate that the marker decreases disease. Additionally, S1_183842019, located in the qTAR_1.2 region (QTL position 1423.60 cM, QTL size 7.37 cM), showed a PVE of 34.1% with an additive effect of -4.54 (Table 2). Furthermore, S1_184245671 in the qTAR_1.3 region (QTL position 1440.38 cM, QTL size 2.84 cM) and S1_184915672 and S1_184914824 in the qTAR_1.4 region (QTL position 1457.50 cM, QTL size 2.43 cM) showed PVEs of 28.8% and 28.0%, with additive effects of -4.24 and -4.18, respectively. Additionally, S1_187588773 was in the qTAR_1.5 region (QTL position 1464.88 cM, QTL size 9.02 cM) and showed a PVE of 20.7% with an additive effect of -4.20 (Table 2). The largest QTL intervals corresponded to the qTAR_1.1, qTAR_1.2 and qTAR_1.5 regions, spanning approximately 12.64 cM, 7.37 cM and 9.02 cM, respectively. The remaining QTLs, qTAR_1.3 and qTAR_1.4, had comparatively smaller intervals of about 2.84 and 2.43 cM, respectively.

**Fig. 7.**
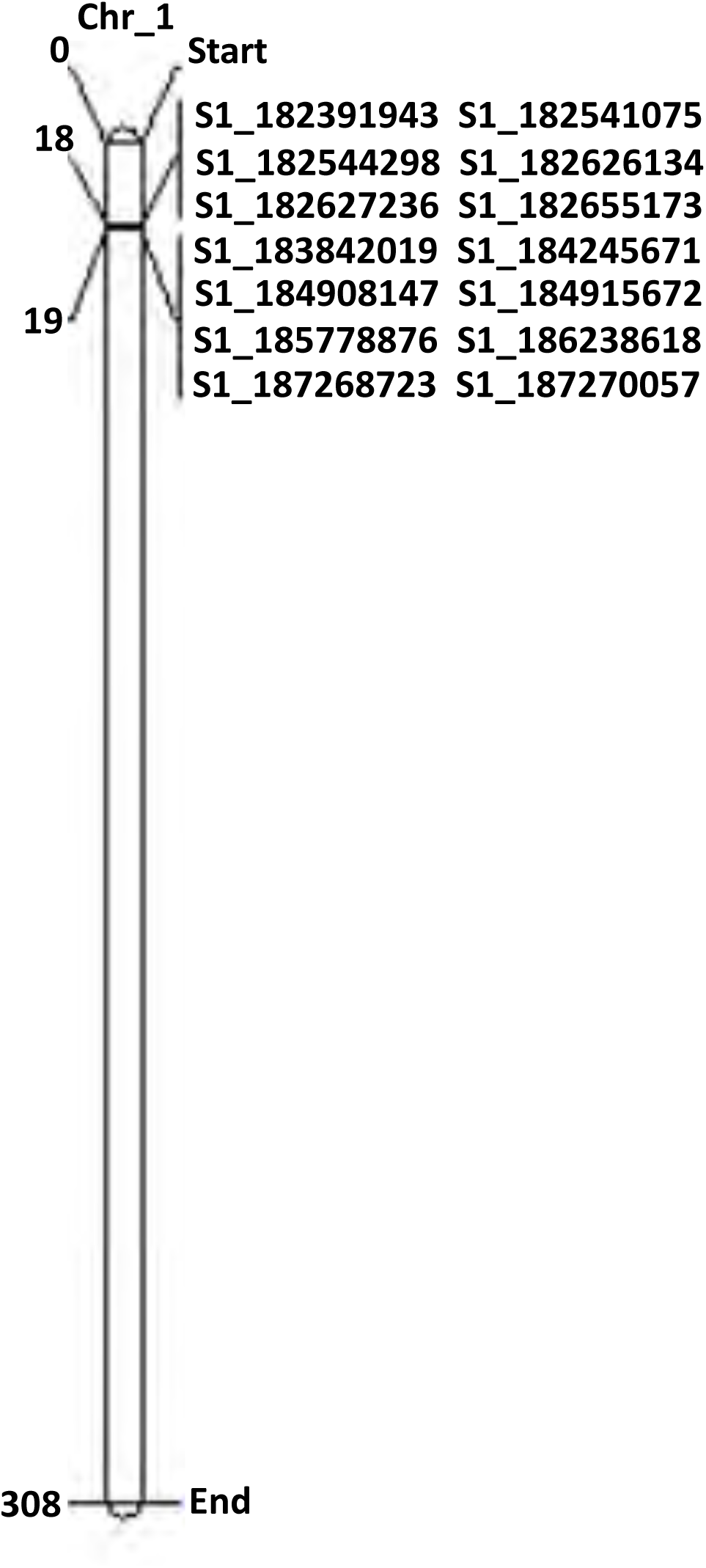
Genetic map of corn chromosome 1, displaying significant SNP markers associated with a Quantitative Trait Locus (QTL) for tar spot resistance using linkage mapping in the Intermated B73 × Mo17 (IBM) maize recombinant-inbred line (RIL) population after two seasons of field testing. SNP markers are scaled according to their physical location (Mb).

**Table 2.** Physical positions and genetic effects of 5 significant Quantitative Trait Loci (QTL) (in bold) and 31 significant Single-Nucleotide Polymorphisms (SNPs) on maize chromosome 1 identified by QTL mapping of tar spot resistance in the Intermated B73 x Mo17 (IBM) recombinant-inbred line (RIL) Syn5 population in Environment I (from 2020) and Environment II (from 2021), where a robust statistical threshold of LOD 3.8, determined by 1,000 permutations (p < 0.05), was used to define significance

|  |  |  |  |  |  | Marker interval<sup>a</sup> |  |  |  |  |
| --- | --- | --- | --- | --- | --- | --- | --- | --- | --- | --- |
| Environment<sup>b</sup> | QTL | LOD | QTL position (cM) | SNP<sup>c</sup> | Marker position (cM) | Ci<sup>d</sup>_low | Ci<sup>d</sup>_high | QTL size (cM) | PVE % (R<sup>2</sup>)<sup>e</sup> | Additive effect<sup>f</sup> |
| I |  | 4.12* |  | S1_182391943<sup>a, c</sup> | 1388.48 |  |  |  | 19.4 | -3.50 |
|  |  | 4.12* |  | S1_182541075<sup>a, c</sup> | 1388.88 |  |  |  | 19.8 | -3.50 |
|  |  | 4.12* |  | S1_182542905<sup>a, c</sup> | 1390.49 |  |  |  | 19.6 | -3.50 |
|  |  | 4.12* |  | S1_182542912<sup>a, c</sup> | 1390.49 |  |  |  | 19.6 | -3.50 |
|  |  | 4.13* |  | S1_182543762<sup>a, c</sup> | 1392.49 |  |  |  | 19.6 | -3.50 |
|  |  | 4.13* |  | S1_182544298<sup>a, c</sup> | 1392.49 |  |  |  | 19.4 | -3.50 |
|  |  | 4.13* |  | S1_182623088<sup>a, c</sup> | 1393.68 |  |  |  | 19.2 | -3.50 |
|  |  | 4.13* |  | S1_182623103<sup>a, c</sup> | 1394.07 |  |  |  | 19.2 | -3.50 |
|  |  | 4.13* |  | S1_182626134<sup>a, c</sup> | 1395.30 |  |  |  | 20.7 | -3.50 |
|  | **qTAR_1.1** | **4.13\*** | **1395.70** | **S1_182627236<sup>a, c</sup>** | **1395.70** | **1386.07** | **1398.53** | **9.46** | **19.4** | **-3.50** |
|  |  | 4.05* |  | S1_182655173<sup>a, c</sup> | 1396.50 |  |  |  | 19.7 | -3.49 |
|  |  | 3.80 |  | S1_182793045<sup>a, c</sup> | 1398.53 |  |  |  | 18.8 | -3.41 |
|  |  | 3.80 |  | S1_182793076<sup>a, c</sup> | 1398.53 |  |  |  | 18.8 | -3.41 |
| I & II |  | 4.27* |  | S1_183811524<sup>a, c</sup> | 1418.68 |  |  |  | 25.1 | -3.64 |
|  |  | 4.27* |  | S1_183811527<sup>a, c</sup> | 1418.68 |  |  |  | 25.1 | -3.64 |
|  | **qTAR_1.2** | **7.43\*\*** | **1423.60** | **S1_183842019<sup>a, c</sup>** | **1423.60** | **1418.68** | **1426.05** | **7.37** | **34.1** | **-4.54** |
|  |  | 7.42** |  | S1_183842131<sup>a, c</sup> | 1424.01 |  |  |  | 34.4 | -4.54 |
|  |  | 7.36** |  | S1_183849097<sup>a, c</sup> | 1425.23 |  |  |  | 34.5 | -4.53 |
|  |  | 7.43** |  | S1_183850352<sup>a, c</sup> | 1426.05 |  |  |  | 29.7 | -4.35 |
| I & II |  | **6.20\*\*** |  | S1_184242660<sup>a, c</sup> | 1439.47 |  |  |  | 28.8 | -4.24 |
|  | **qTAR_1.3** | **6.38\*\*** | **1440.27** | **S1_184245671<sup>a, c</sup>** | **1440.27** | **1439.47** | **1442.31** | **2.84** | **28.9** | **-4.32** |
|  |  | 5.98** |  | S1_14623957<sup>a, c</sup> | 1442.31 |  |  |  | 29.1 | -4.19 |
| I |  | 6.14** |  | S1_184908147<sup>a, c</sup> | 1455.87 |  |  |  | 28.0 | -4.18 |
|  |  | 6.46** |  | S1_184914824<sup>a, c</sup> | 1457.09 |  |  |  | 31.7 | -4.30 |
|  | **qTAR_1.4** | **6.55\*\*** | **1457.50** | **S1_184915672<sup>a, c</sup>** | **1457.50** | **1455.87** | **1458.30** | **2.43** | **29.9** | **-4.33** |
|  |  | 6.18** |  | S1_185778876<sup>a, c</sup> | 1458.30 |  |  |  | 28.1 | -4.20 |
| II |  | 3.89* |  | S1_186238573<sup>a, c</sup> | 1459.52 |  |  |  | 19.8 | -4.13 |
|  |  | 3.89* |  | S1_186238606<sup>a, c</sup> | 1462.01 |  |  |  | 19.8 | -4.13 |
|  |  | 4.15* |  | S1_187268723<sup>a, c</sup> | 1463.66 |  |  |  | 20.1 | -4.21 |
|  |  | 4.17* |  | S1_187270057<sup>a, c</sup> | 1464.47 |  |  |  | 20.9 | -4.20 |
|  | **qTAR_1.5** | **4.18\*** | **1464.88** | **S1_187588773<sup>a, c</sup>** | **1464.88** | **1459.52** | **1468.54** | **9.02** | **20.7** | **-4.20** |

<sup>a</sup> 95% probability that the indicated QTL is between these positions on the B73 RefGen\_v4 genome sequence.
<sup>b</sup> Environment: I (2020) and II (2021), at the Pinney Purdue Agricultural Center, Wanatah, IN.
<sup>c</sup> Single-nucleotide polymorphism marker flanking the QTL with LOD score of $\geq 3.8$ .
<sup>d</sup> Ci is the confidence interval of the QTL locus.
<sup>e</sup> Phenotypic variance of a single QTL.
<sup>f</sup> Additive effects of the QTL. Negative additive effects indicate that the QTL reduces disease severity.
Significance levels for LOD scores: $p = 0.1$ (all values), $p = 0.05$ (\*), and $p = 0.01$ (\*\*).

qTAR_1.2, qTAR_1.3 and qTAR_1.4 emerged as the strongest QTLs in our analysis, evidenced by their statistical significance and LOD scores. The S1_183842019 marker, located at the qTAR_1.2 peak, exhibited the highest LOD score of 7.43 and the highest PVE (34.1%) among all detected loci. Other markers, including S1_184245671 and S1_184915672 located at the qTAR_1.3 and qTAR_1.4 peaks, exhibited LOD scores of 6.38 and 6.55, explaining 28.9% and 29.9% of the phenotypic variance, respectively. Overall, the identification of a consistent, major QTL on chromosome 1 across twoyears provides valuable information for understanding the complex genetic basis of tar spot resistance in corn. These QTLs represent novel findings, as they have not been reported previously for tar spot resistance.

### Identification of candidate genes associated with tar spot resistance

Candidate genes potentially involved in tar spot resistance were identified by mapping QTL intervals to the maize B73 reference genome (V4). The analysis of five distinct genomic regions on chromosome 1, designated qTAR_1.1 through qTAR_1.5, revealed 74 putative genes within the interval from 184.2 Mb to 189.6 Mb (Table 3). Of the 74 genes, eight high-priority candidates were identified as residing directly at the SNP peak positions. In qTAR_1.1, critical defense signaling and regulatory genes were co-localized precisely with the peak markers, including a bZIP transcription factor (Zm00001d031272), a RING/U-box superfamily protein (Zm00001d031273), and a Chaperone protein htpG (Zm00001d031274) (Table 3). These genes are known to mediate E3 ubiquitin ligase activity and maintain cellular stability and are essential components of pathogen-associated molecular pattern (PAMP)-triggered immunity. Further along the cluster, the peak SNPs in QTL qTAR_1.2 and qTAR_1.3 were associated with Endomembrane protein 70 (Zm00001d031317) and a DNA-directed RNA polymerase (Zm00001d031333), suggesting a possible role for intracellular transport and transcriptional reprogramming in the resistance response. Near the qTAR_1.5 peak, a C2H2-type zinc finger protein (Zm00001d031416) and a frataxin homolog (Zm00001d031418) were identified, highlighting the potential importance of transcriptional regulation and mitochondrial homeostasis to resistance. Additionally, an unannotated gene (Zm00001d031352) was located directly on major peak qTAR_1.4, representing a potential novel locus for further functional characterization.

**Table 3.**
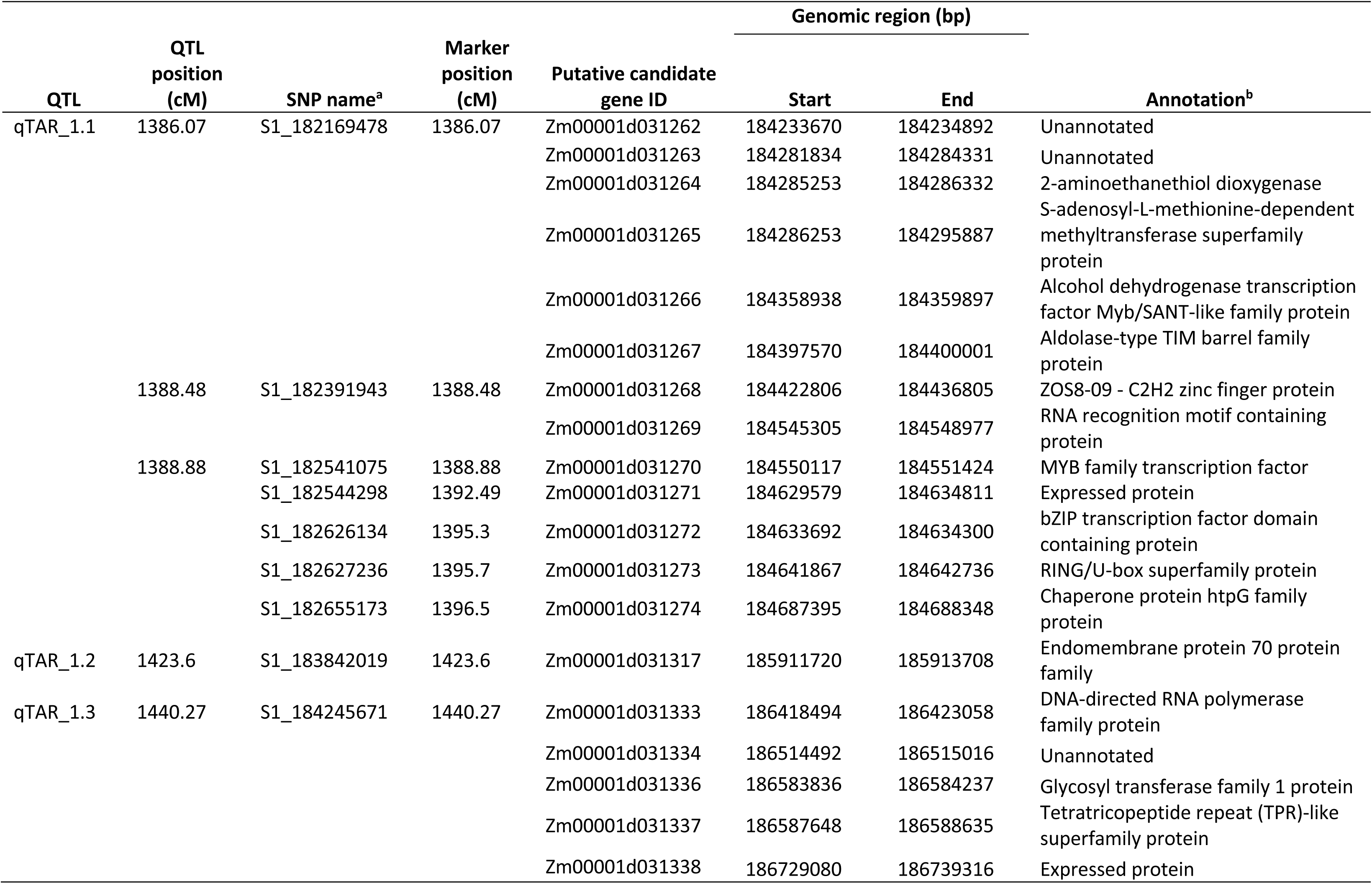
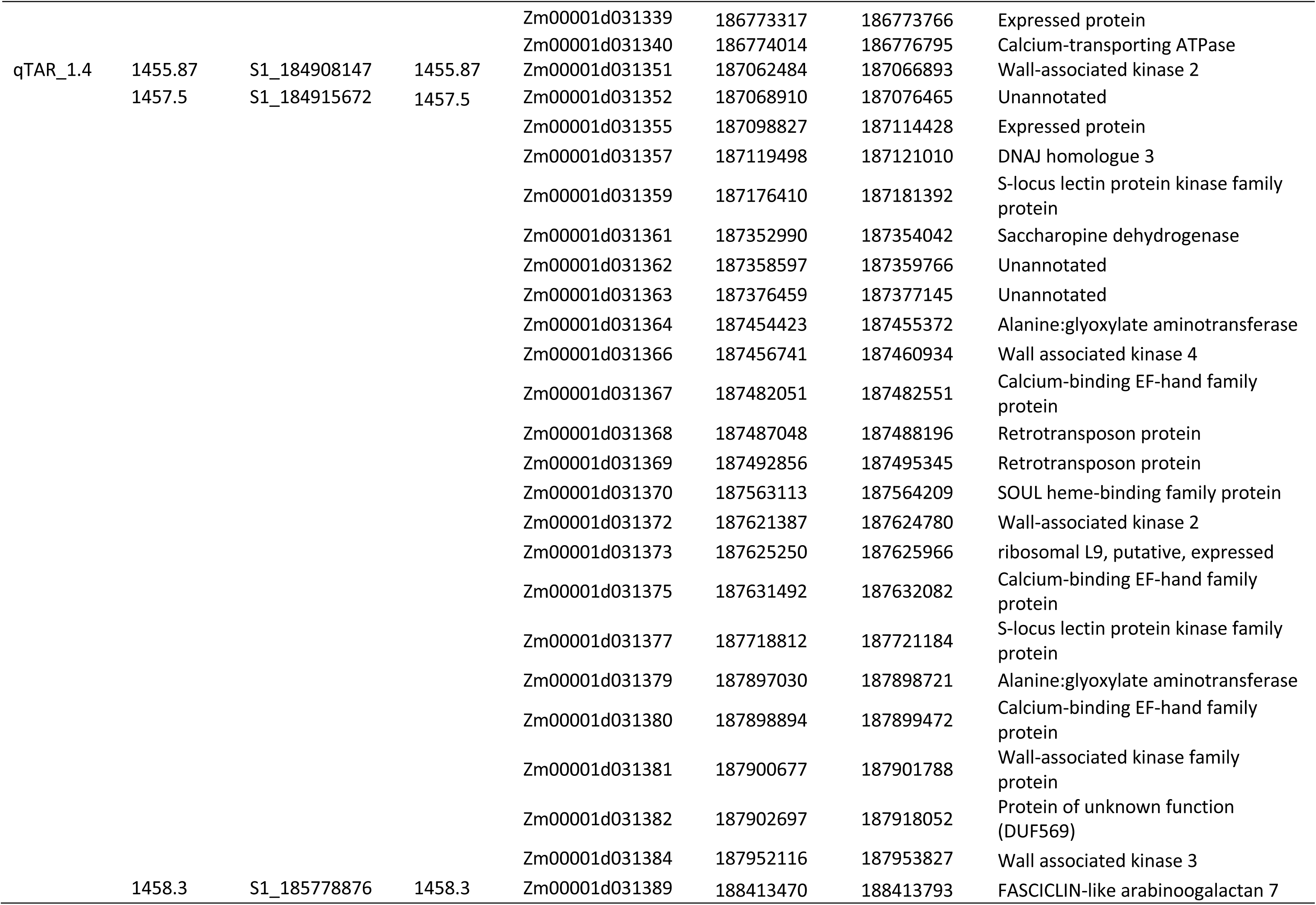
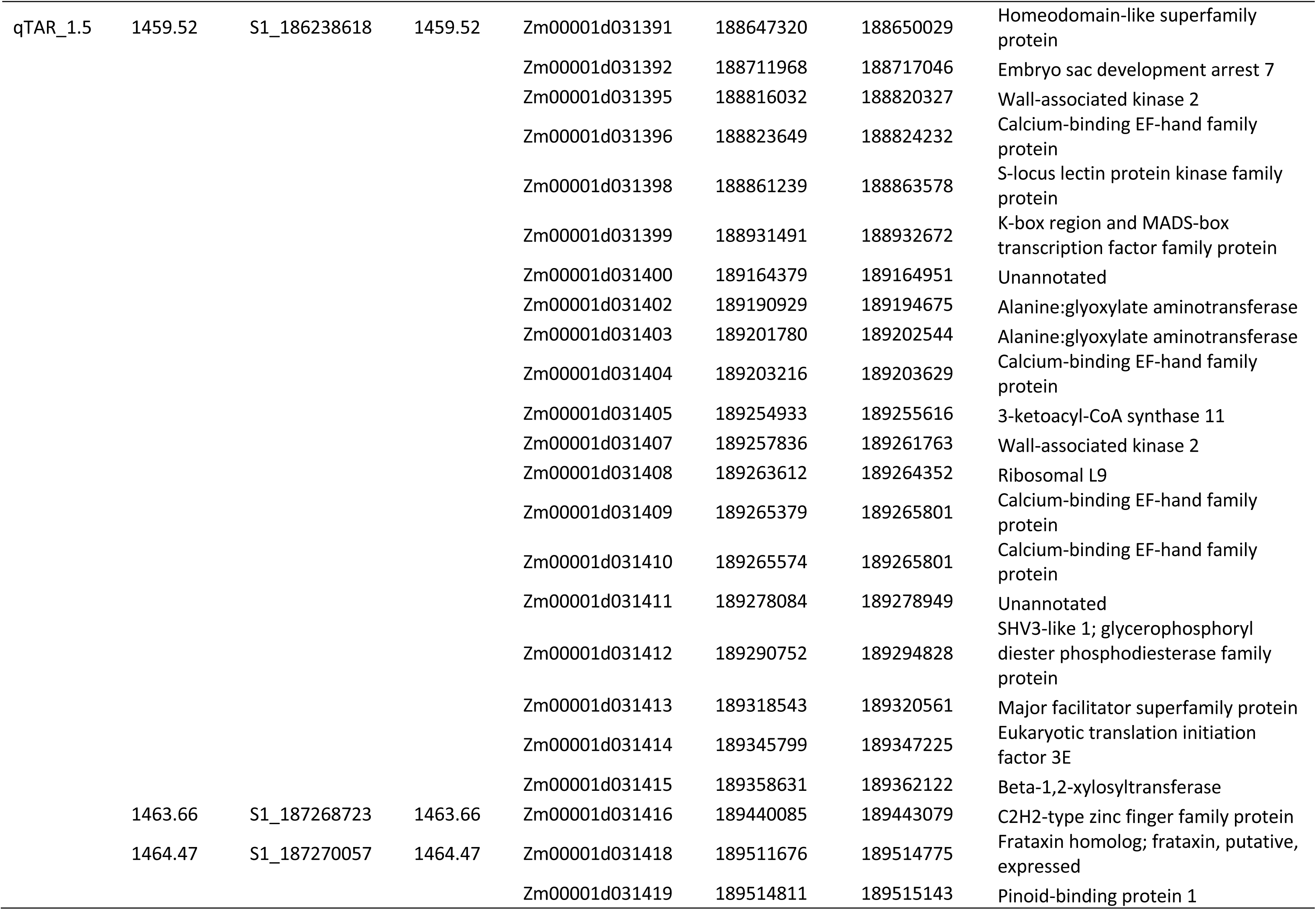
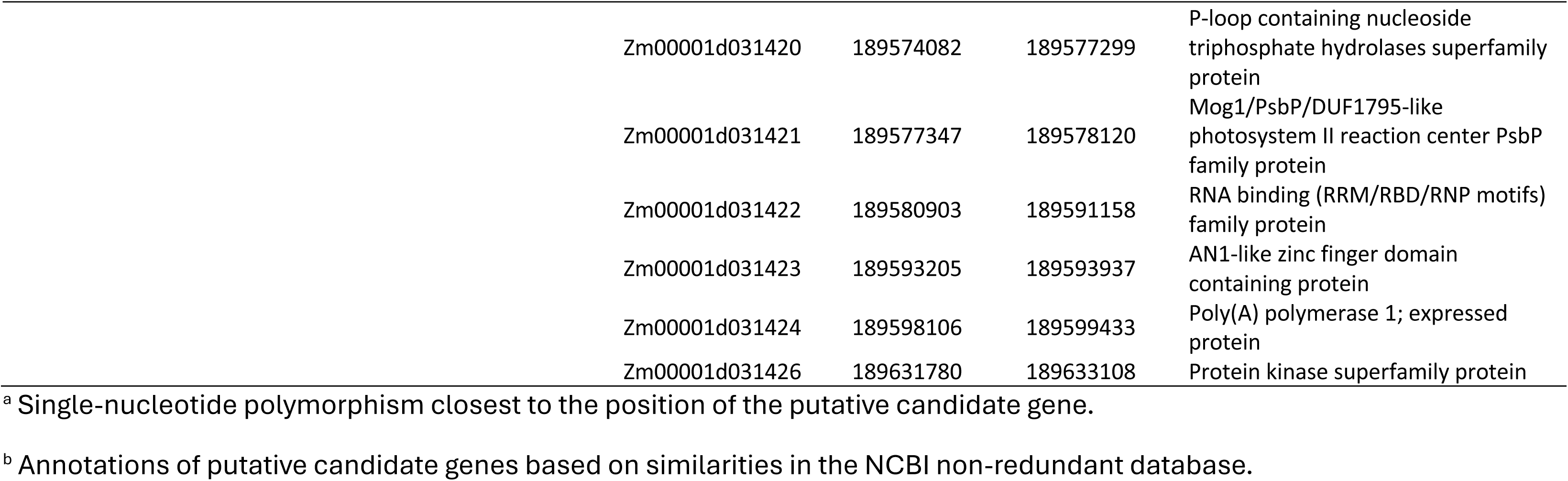
List of candidate genes associated with tar spot resistance in the recombinant-inbred lines of the Intermated B73 × Mo17 (IBM) maize recombinant-inbred line (RIL) Syn5 population.

Beyond the peak-associated genes, the broader intervals contained several gene families that are considered essential for plant defense. qTAR_1.1 included MYB family transcription factors (Zm00001d031266, Zm00001d031270), which regulate the phenylpropanoid pathway. The qTAR_1.3 region was enriched with enzymes for secondary metabolism, such as glycosyltransferase (Zm00001d031336) and multiple alanine:glyoxylate aminotransferases, which provide biochemical fortification against fungal penetration. Notably, a significant concentration of Wall-Associated Kinases (WAKs) (6 genes) and S-locus lectin protein kinases (3 genes) was found across qTAR_1.4 and qTAR_1.5, which included 24 and 29 genes in total, respectively. These included multiple copies of WAK 2, WAK 3, and WAK 4, which serve as primary receptors monitoring cell wall integrity. These regions also harbor numerous Calcium-binding EF-hand family proteins, that are critical for early-stage signal transduction following pathogen recognition.

Overall, the dense physical clustering of these 74 genes within a 5-Mb interval, coupled with the immediate proximity of eight high-priority candidates to highly significant SNPs underscores the potential importance of these genes to the resistance response. Characterizing these specific genes is vital for understanding the complex genetic architecture of tar spot resistance and identifying the primary drivers of the host immune response in maize.

## Discussion

The rapid spread of tar spot across North America from its indigenous home in South and Central America has established *P. maydis* as a primary threat to maize production. The substantial yield reductions observed in recent years underscore the urgency of shifting from reactive management to proactive genetic resistance. However, the phenotypic expression of tar spot is not a simple binary trait; instead, it is a product of complex genotype and environment interactions. Our results highlight that precise phenotypic assessment in environments with higher disease pressure is the only reliable way to distinguish true genetic resistance from environmental noise. By utilizing the IBM RIL population over two years we were able to filter out these variables and pinpoint stable resistance QTL.

The evaluation of the Intermated B73 x Mo17 (IBM) RIL population over two growing environments (2020 and 2021) revealed substantial phenotypic variation. Use of the IBM-94 core subset as a strategic starting point allowed for an efficient survey of this genetic diversity while optimizing field resources. This approach served as a successful proof of concept, validating that the phenotypic variation within this representative subset of the total IBM population was robust enough to allow genetic mapping without immediately requiring the full, larger population.

Although conducted at the same general field location, these two years were classified as distinct environments due to variations in temperature, rainfall and humidity (Supplemental Table S1). Our findings consistently validate B73 as a moderately resistant parent and Mo17 as a highly susceptible parent, aligning with previously published data (Singh et al. 2023). B73 exhibited significantly lower disease severity in both Environment I (8.3%) and Environment II (10.4%) compared to Mo17 (16.9% and 33.4%, respectively), confirming stable resistance under varying disease pressure (Supplemental Table S2). While B73 consistently exhibited moderate resistance and Mo17 displayed susceptibility, the shift from a positively skewed distribution in 2020 to a multimodal distribution in 2021 highlights the dynamic nature of tar spot epidemiology (Fig. 1). The increased disease pressure in Environment II (2021), where mean disease severity (MDS) ranged from 10 to 35%, likely resulted from weather conditions favorable to the pathogen such as high humidity and prolonged leaf wetness and a diverse inoculum source (Supplemental Table S2). These factors are well-documented drivers of tar spot development (Hock et al. 1995; Groves et al. 2020; Kleczewski et al. 2019; Webster et al. 2023). This increased pressure stretched the phenotypic range, allowing for a clearer differentiation between truly resistant RILs and those that merely escaped infection in the lower-pressure environment of 2020.

The continuous distribution of AUDPC values observed in both years is characteristic of a quantitative trait controlled by multiple small-effect QTLs (Fig. 3B). The presence of lines at both phenotypic extremes of the AUDPC scale in both years indicates that the IBM population is an effective tool for mapping the genetic architecture of *P. maydis* resistance. Despite the overall increase in disease in 2021, the jitter points in the box plot (Fig. 3B) show that certain RILs maintained relatively low AUDPC values across both environments. These consistently resistant lines are of particular interest for further breeding efforts, as they likely harbor stable alleles that are less sensitive to environmental fluctuations. Conversely, the increased spread of AUDPC scores in 2021 allowed for better differentiation of moderately susceptible lines that appeared resistant under the lighter disease pressure of 2020.

A critical finding of this study is the high repeatability of the disease scores across environments. Despite environmental shifts, Pearson correlation analysis showed a strong, highly significant positive correlation (r = 0.8706, p < 0.0001). The coefficient of determination (R^2^ = 0.7579) indicates that approximately 76% of the phenotypic variance is attributed to the genetic background rather than environmental noise (Fig. 2). Standardized checks validated the phenotyping protocol, with resistant RILs consistently maintaining low severity and susceptible genotypes occupying the high-severity quadrant of the regression (Fig. 2). RILs that consistently fell into the low-severity quadrant across both years represent high-value germplasm for introgression into elite breeding lines. While outliers like RIL #M0043 highlight specific gene-by-environment interactions, the overarching trend suggests that the resistance identified here is durable and may be suitable for deployment across diverse environments in North America.

The ANOVA results further support these findings, showing that variation among genotypes was the primary driver of the phenotypic differences observed (p < 0.001) (Table. 1). This confirms that the resistance observed in B73 is not only superior to Mo17 but also is segregating successfully in the RIL population in a predictable manner. The low residual mean square (10.7) further validates the experimental precision of the disease severity protocol used for scoring. This level of repeatability is essential for breeding programs, suggesting that selections made in one environment will likely maintain resistance across varied field conditions.

A central finding of this study is the identification of a consistent, major cluster of five significant QTL regions (qTAR_1.1 to qTAR_1.5) for tar spot on corn chromosome 1. These regions exceeded the genome-wide threshold for statistical significance (LOD 3.8) in both environments, marking them as stable genetic drivers of the resistance (Fig. 3A and B, Table 2). Minor peaks on chromosomes 8 and 10 were higher than LOD 2.0 in both years but failed to meet the stringent LOD 3.8 threshold for statistical significance overall (Fig. 3A and B). Interestingly, while previous studies in CIMMYT and DTMA germplasm identified resistance on chromosome 8 (Mahuku et al. 2016; Cao et al. 2017; Ren et al. 2022; Yan et al. 2022), our results suggest a different genetic architecture in the IBM population. The minor peaks identified on chromosome 8 in our analyses remained below the genome-wide significance threshold, suggesting that the tar spot resistance in B73 is driven by a novel locus on chromosome 1 that has not been reported previously, rather than qRtsc8-1, which was identified in tropical maize germplasm. We provisionally call this new resistance QTL qRtsc1-1.

Previously, QTLs on chromosome 1 have been associated with resistance to various maize diseases including Northern Leaf blight (NLB), Stewart’s Wilt, Southern Leaf Blight (SLB), Grey Leaf Spot (GLS), Common Rust, and Maize Streak Virus, suggesting potentially shared genetic mechanisms or closely linked resistance clusters (Jamann et al. 2014; Wisser et al. 2006, Zwonitzer et al. 2010, Nair et al. 2015, Martins et al. 2019). Combining qRtst1-1 with the previously identified qRtst8-1 will likely result in a greater level of tar spot resistance than either QTL alone.

The linkage mapping results reveal a highly localized yet possibly complex genetic architecture for tar spot resistance on chromosome 1. One possible explanation for the 31 significant SNPs across five regions (qTAR_1.1to qTAR_1.5) may be that resistance in B73 may be governed by multiple, tightly linked loci (Table 3). Specifically, qTAR_1.2, qTAR_1.3, and qTAR_1.4 emerged as the most statistically significant regions. The marker S1_183842019, located within the qTAR_1.2 peak, exhibited the highest LOD score (7.43) and explained the greatest portion of phenotypic variance (34.1%). The negative additive effects observed across the SNPs (ranging from -3.50 to -4.54) indicate that alleles from the resistant parent effectively reduce disease severity (Table 2). Other significant markers on chromosome 1 (e.g., S1_184245671 in qTAR_1.3, and S1_184915672 in qTAR_1.4) had LOD scores exceeding 6.0, further indicating their individual possible contributions to resistance (Table 2).

The variation in QTL interval sizes offers insights into the genomic resolution of this study. The relatively smaller intervals for qTAR_1.3 (2.84 cM) and qTAR_1.4 (2.43 cM) suggest that these loci are well defined, potentially narrowing the search for candidate genes in future fine-mapping efforts. In contrast, the larger spans of qTAR_1.1 (12.64 cM) and qTAR_1.5 (9.02 cM) may harbor multiple genes or represent regions with lower recombination frequencies in the IBM population. A major highlight of these findings is their novelty; none of the five identified regions have been previously associated with tar spot resistance in other mapping studies. While previous literature has often focused on chromosome 8 for tar spot resistance, the discovery of this consistent, major genetic driver on chromosome 1 provides a new resource for corn improvement.

Our linkage mapping analysis successfully narrowed the genomic regions associated with tar spot resistance on chromosome 1, enabling the in-depth identification of potential candidate genes within these QTL intervals. By precisely mapping the significant SNP peaks to the well-annotated B73 reference genome (V4), we identified 74 candidate genes across five significant SNP-associated QTL regions (qTAR_1.1 to qTAR_1.5). Eight high-priority candidate genes reside directly at the SNP peak positions, suggesting a strong association with the resistance phenotype (Table 3).

A striking feature of the qTAR_1.4 and qTAR_1.5 intervals is the high density of Wall-Associated Kinases (WAKs, WAK 2, 3, and 4). This mirrors the architecture of known resistance genes such as *Htn1*, which also utilizes WAKs to confer resistance against leaf pathogens (Zuo et al. 2015). These receptors monitor cell wall integrity and are known to trigger defense responses, such as an oxidative burst, upon sensing fungal penetration (Yang et al. 2017). The proximity of these kinases to highly significant SNPs, such as S1_184915672 and S1_187268723, reinforces the hypothesis that WAK-mediated recognition may be a central pillar of the resistance phenotype in this population (Table 3). The co-localization of WAKs with numerous calcium-binding EF-hand family proteins suggests a possible early-stage signal transduction hub that may convert the physical recognition of the pathogen into biochemical signals (Boudsocq and Sheen, 2013).

The eight high-priority candidates situated at the SNP peaks provide a compelling narrative for possible transcriptional and post-translational control of the resistance response. The bZIP (Zm00001d031272) and C2H2-type zinc finger (Zm00001d031416) transcription factors, alongside MYB proteins, may serve as early responders that modulate the expression of stress-responsive genes and the phenylpropanoid pathway in maize (Du et al. 2012). Furthermore, the possible post-translational control provided by the RING/U-box superfamily protein (Zm00001d031273), an E3 ubiquitin ligase, represents a critical component of pathogen-associated molecular pattern (PAMP)-triggered immunity (PTI) (Yee and Goring, 2009; Trujillo and Shirasu, 2010).

Beyond signaling, the plant may actively maintain cellular homeostasis during the stress of infection. The presence of chaperone protein htpG (Zm00001d031274) and endomembrane protein 70 (Zm00001d031317) suggests that maintaining protein folding and intracellular transport may be vital for sustained defense. Heat shock proteins and chaperones like htpG are essential for the maturation and stability of R proteins and other signaling complexes during biotic stress responses (Kadota and Shirasu, 2012). Simultaneously, the enrichment of enzymes such as glycosyltransferases and alanine:glyoxylate aminotransferases suggests a metabolic shift towards biochemical fortification (Umar et al. 2021). These enzymes contribute to the synthesis of secondary metabolites that provide a physical and chemical barrier against fungal hyphae (Vance et al. 1980; Dixon and Paiva, 1995). Interestingly, the frataxin homolog (Zm00001d031418) may play a role in maintaining mitochondrial homeostasis during this high-energy metabolic shift (Busi et al. 2006). The identification of an unannotated gene (Zm00001d031352) directly on a major peak further underscores the potential for discovering novel elements of resistance to tar spot in maize.

Overall, the candidate genes identified in this study provide a rich resource for dissecting the genetic basis of tar spot resistance. Our results confirm that the IBM-94 core subset is highly effective for initial genetic screens, providing sufficient mapping resolution to identify critical genomic regions while optimizing experimental resources. The enrichment of genes involved in signaling, transcription, cell wall integrity, and metabolism strongly support their potential role in resistance. However, to further refine their genomic intervals and confirm the stability of the identified QTLs, future research should expand this analysis to the full IBM and other mapping populations. Such efforts, combined with functional characterization via CRISPR-Cas9 or by testing existing mutant lines for the candidate genes in B73 will ultimately elucidate the molecular mechanisms of resistance and accelerate the development of tar spot-resistant cultivars through targeted breeding strategies.

## Acknowledgments

We thank Torbert Rocheford at Purdue University for providing seeds of the IBM parental lines and segregating progeny, and Wily Rodrigo Sic-Hernandez, Sandra Victoria Gomez-Gutierrez, Kristy Lin, Brett Kang, and J. Ravellette at Purdue University plus Ian Thompson at USDA-ARS for assistance with field trial establishment and maintenance.

## Supplementary Figures

**Supplementary Fig. S1.** Genome-wide linkage analysis of tar spot resistance in the Intermated B73 × Mo17 (IBM) maize recombinant-inbred line (RIL) population evaluated at the early disease development stage during the 2020 field season. Logarithm of the odds (LOD) profiles across 10 chromosomes are shown for 92 IBM RILs evaluated in the 2020 field environment. The x axis represents the genetic position across all 10 chromosomes, and the y axis indicates the LOD score. No significant quantitative trait loci (QTL) were detected, as all peaks remained below the significance threshold of LOD 3.8 (p < 0.05), determined through 1,000 permutations.

**Supplementary Fig. S2.** Logarithm of the odds (LOD) profiles for tar spot resistance on maize chromosome 1. Quantitative trait loci (QTL) were identified using the Intermated B73 × Mo17 (IBM) Syn5 recombinant inbred line (RIL) population. The horizontal, red, dashed line indicates the genome-wide significance threshold of LOD 3.8 (p < 0.05), established via 1,000 permutations. The x axis represents the genetic distance in centimorgans (cM), and the y axis represents the LOD score. (**A**) LOD profile for tar spot response based on mean disease severity data from the 2020 growing season. (**B**) LOD profile for tar spot response based on mean disease severity data from the 2021 growing season.

## Supplementary Tables

**Supplementary Table S1.** Weather data for the Pinney Purdue Agriculture Centre (PPAC), Wanatah, IN for the periods of 5/1/2020 to 11/4/2020 and 5/1/2021 to 11/5/2021.

**Supplementary Table S2.** Tar spot disease severity scores for 92 Intermated B73 × Mo17 (IBM) recombinant-inbred lines (RILs) (seeds from two of the 94 RILS did not germinate) including parents B73 and Mo17 over two growing seasons: the 2020 field environment (Environment I) and the 2021 field environment (Environment II) at the Pinney Purdue Agriculture Center (PPAC) in Wanatah, IN.

**Supplementary Table S3.** Phenotypic correlation and stability of tar spot severity in 92 recombinant-inbred lines (RILs) from the Intermated B73 × Mo17 (IBM) population (seeds from two of the 94 RILS did not germinate). Comparative analysis of mean disease severity (%) for 92 RILs and parental checks (B73 and Mo17) across two environments (2020 and 2021).3

