## Supplemental Fig. S2 for "Mapping and Genetic Dissection of Novel Tar Spot Resistance QTL on Maize Chromosome 1"

### Slide 1
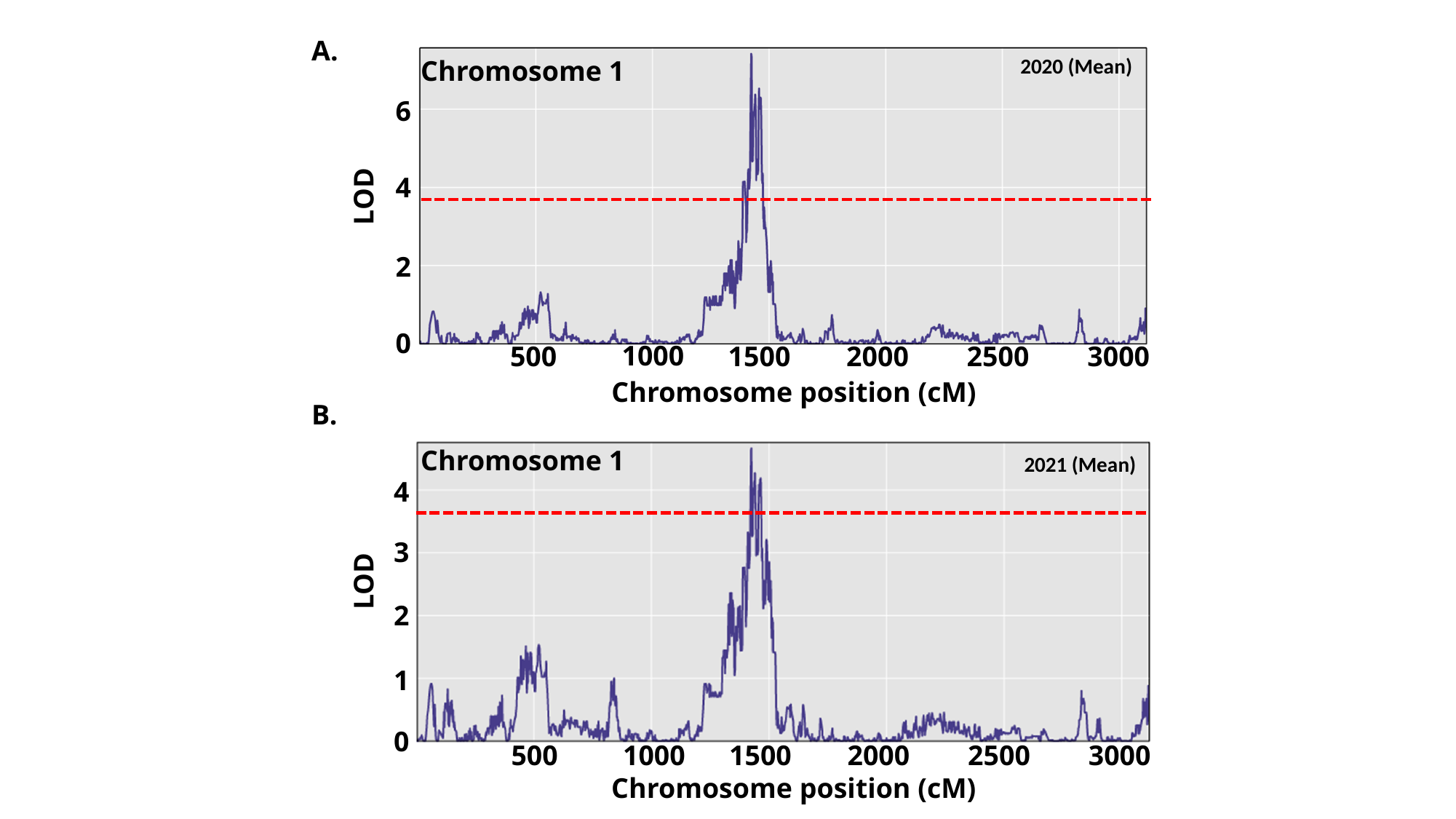

A.
 2020 (Mean)
Chromosome 1
6
4
LOD
2
0
1000
500
1500
2000
2500
3000
Chromosome position (cM)
B.
Chromosome 1
 2021 (Mean)
4
3
LOD
2
1
0
500
1000
1500
2000
2500
3000
Chromosome position (cM)

### Slide 2
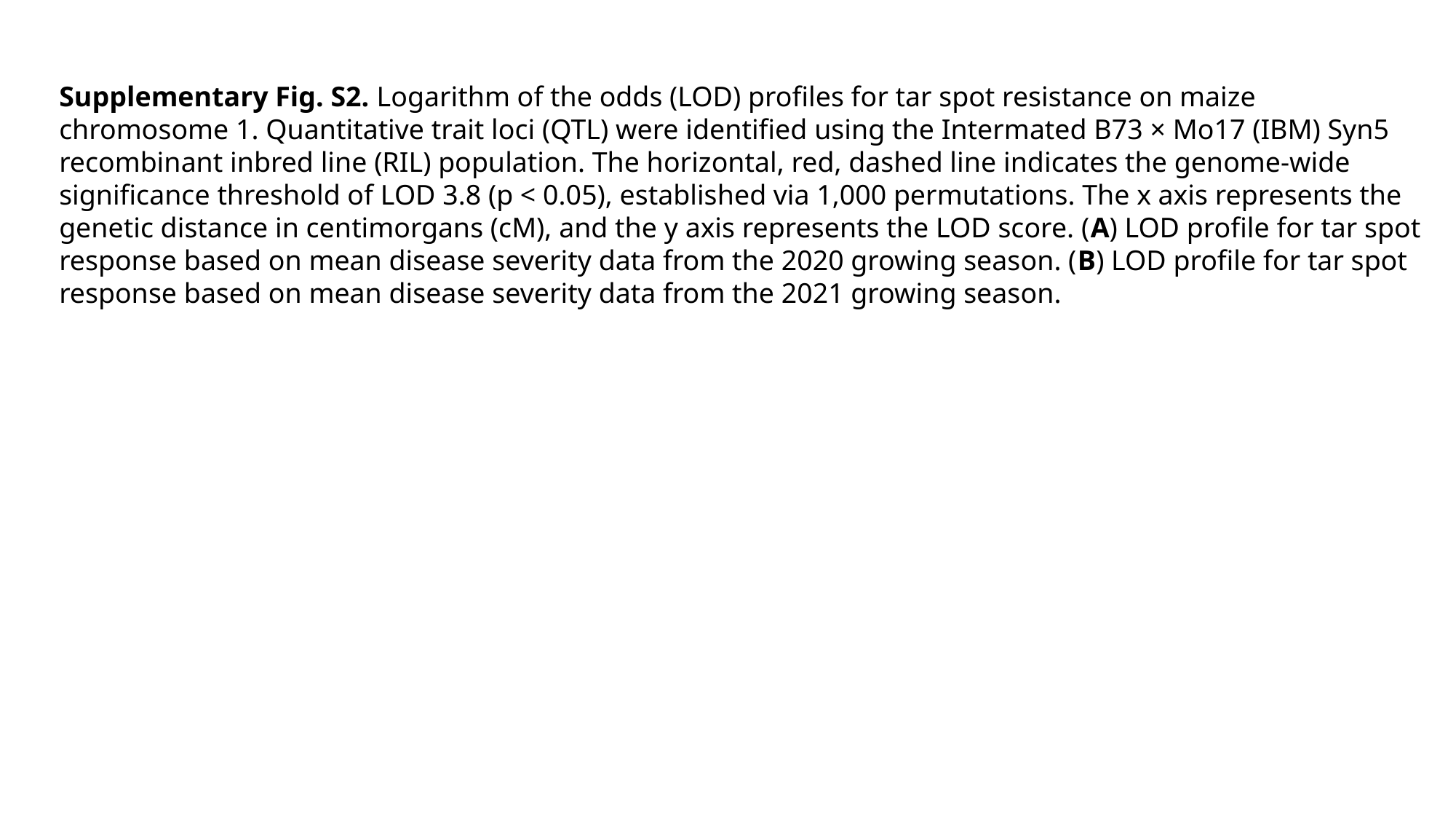

Supplementary Fig. S2. Logarithm of the odds (LOD) profiles for tar spot resistance on maize chromosome 1. Quantitative trait loci (QTL) were identified using the Intermated B73 × Mo17 (IBM) Syn5 recombinant inbred line (RIL) population. The horizontal, red, dashed line indicates the genome-wide significance threshold of LOD 3.8 (p < 0.05), established via 1,000 permutations. The x axis represents the genetic distance in centimorgans (cM), and the y axis represents the LOD score. (A) LOD profile for tar spot response based on mean disease severity data from the 2020 growing season. (B) LOD profile for tar spot response based on mean disease severity data from the 2021 growing season.
